# Transcriptional expression changes during compensatory plasticity in the prothoracic ganglion of the adult cricket *Gryllus bimaculatus*

**DOI:** 10.1101/2021.11.24.469824

**Authors:** Felicia Wang, Harrison Fisher, Maeve Morse, Lisa L. Ledwidge, Jack O’Brien, Sarah E. Kingston, Justin Beckman, Jasmine J. Johnson, Lyn S. Miranda Portillo, Tabarak N. Al Musawi, Alexandra W. Rubenstein, David A. Michaelson, Hadley Wilson Horch

**Affiliations:** Department of Biology, Bowdoin College, 6500 College Station, Brunswick, Maine 04672 USA.

**Keywords:** RNASeq, Transcriptome, Dendritic Plasticity, GO term analysis, Differential expression, Injury

## Abstract

Most adult organisms are limited in their capacity to recover from neurological damage. The auditory system of the Mediterranean field cricket, *Gryllus bimaculatus*, presents a compelling model for investigating neuroplasticity due to its unusual capabilities for structural reorganization into adulthood. Specifically, the dendrites of the central auditory neurons of the prothoracic ganglion sprout in response to the loss of auditory afferents. Deafferented auditory dendrites grow across the midline, a boundary they normally respect, and form functional synapses with the contralateral auditory afferents, restoring tuning-curve specificity. The molecular pathways underlying these changes are entirely unknown. Here, we used a multiple k-mer approach to re-assemble a previously reported prothoracic ganglion transcriptome that included ganglia collected one, three, and seven days after unilateral deafferentation in adult, male animals. We used EdgeR and DESeq2 to perform differential expression analysis and we examined Gene Ontologies to further understand the potential molecular basis of this compensatory anatomical plasticity. Enriched GO terms included those related to protein translation and degradation, enzymatic activity, and Toll signaling. Extracellular space GO terms were also enriched and included the upregulation of several protein yellow family members one day after deafferentation. Investigation of these regulated GO terms help to provide a broader understanding of the types of pathways that might be involved in this compensatory growth and can be used to design hypotheses around identified molecular mechanisms that may be involved in this unique example of adult structural plasticity.

## Background

Most adult organisms, especially mammals, are limited in their capacity to adapt and recover from neurological damage (1, 2). The Mediterranean field cricket, *Gryllus bimaculatus*, provides a model of neuroplasticity due to its demonstrated ability to compensate for neuronal damage with novel dendritic growth and synapse formation, even into adulthood. Specifically, the central auditory system, much of which resides in the prothoracic ganglion, reorganizes in response to deafferentation caused by unilateral transection of auditory afferents in the adult (3–5).

In *G. bimaculatus*, auditory information is transduced by the auditory organs, located on the prothoracic limbs. Auditory afferents receive the sensory stimuli and convey this information into the prothoracic ganglion where they form synapses with several different auditory neurons (6, 7). These neurons exist as mirror image pairs and their dendritic arbors remain localized ipsilateral to the auditory input, typically not projecting contralaterally across the midline (8). However, previous research has shown that after amputation of the prothoracic leg in the adult, which removes the auditory organ and severs the afferents, the deafferented dendrites of the ipsilateral auditory neurons sprout across the midline and form functional synapses with the intact auditory afferents on the contralateral side. This reorganization is evident whether deafferentation occurs in larvae (9, 10) or adults (3–5). Various aspects of the physiological consequences of this compensatory behavior have been studied (3,9,10), however little is known about the molecular pathways and mechanisms underlying this growth.

Although the genome has only just become publicly available (11), various *de novo* transcriptomes have been created for use in this species (12–14). Recently, a *de novo* transcriptome of the prothoracic ganglion was assembled in an attempt to understand the molecular basis of the compensatory response (15). This transcriptome was built with RNA from individual prothoracic ganglia of both control and deafferented adult male crickets. Initially, this transcriptome was assembled and mined for the presence of developmental guidance molecules. These guidance molecules are known to play a well-conserved role in regulating the specific growth of axonal and dendritic projections during the development of many species (16, 17). While these molecules have mainly been studied for their role in development, it has also been suggested that alterations in their expression may influence the ability of axons and dendrites to recover from injury in adulthood (15,18–20). Mining this cricket transcriptome revealed that many well-conserved developmental guidance molecules, including slit, ephrins, netrins, and semaphorins, were present within the adult prothoracic transcriptome (15). However, it is still unknown whether the expression of these transcripts, or any other transcripts, are significantly altered during this compensatory growth process.

The goal of this study was to better understand the underlying molecular control of the compensatory growth behavior observed in the cricket. We assembled a new, more representative and less redundant transcriptome of the cricket prothoracic ganglion using multiple k-mer values during the assembly process. We also utilized the Evidential Gene *tr2aacdsmRNA* classifier to reduce redundancies (21). This new transcriptome was used to analyze changes in expression levels one, three, and seven days post-deafferentation. The identified genes were then analyzed using GO annotation analysis to determine the classes of genes that are differentially regulated over the course of the injury response. By performing this analysis, it was possible to discover changes in gene expression that occur during the compensatory growth response, allowing for insights into possible pathways or key molecules critical to this process.

## Results and Discussion

### Transcriptome Assembly and Analysis

This transcriptomic study focused on the cricket, *Gryllus bimaculatus*, whose nervous system has been shown to have an unusual level of adult structural plasticity (3–5). We deafferented sensory neurons, including the auditory neurons, in the prothoracic ganglion of the adult cricket, by unilateral amputation of the prothoracic leg at the femur. Control amputations consisted of removal of the distal tip of the tarsus. We harvested prothoracic ganglia one, three, and seven days post-amputation. These time points were designed to capture transcriptional changes in response to the injury (one day), during initial sprouting (one and three days), growth across the midline (three and seven days), and *de novo* synapse formation (3, 22).

Although a *G. bimaculatus* prothoracic ganglion transcriptome from this tissue was previously assembled, analysed, and mined (15), the present study re-assembled a new transcriptome based on those original cleaned and trimmed RNA-Seq reads. Five individual *de novo* transcriptomes were built in Trinity using five different k-mer lengths (21, 25, 27, 30, and 32). Transcriptome construction with longer k-mer lengths produces more reliable contigs, though biased toward highly expressed transcripts. In a complementary fashion, using a shorter k-mer length produces a more exhaustive set of contigs though also one more prone to noise (23, 24). This trade-off between bias and noise induced by the choice of k-mer length suggests how a higher quality *de novo* assembly can be derived by integrating multiple k-mer lengths into an analysis (23). By combining results across k-mer lengths, we ensured that a complementary selection of contigs was included in the analysis (23,25,26). Correspondingly, we combined the five assemblies into a single reference transcriptome and subsequently filtered redundancies and fragments.

The individual assemblies had an N50 ranging from 1,219 - 2,341, with the longer k-mer assemblies yielding a longer N50 (Table 1). The median, average, and maximum contig length also increased as the k-mer length was increased. The total number of Trinity “genes” ranged from 283,278 to 351,829, with higher k-mer assemblies yielding fewer predicted genes. The GC content for each assembly remained roughly constant, between 40-41%. The overall alignment was greater than 98%, with multimapping percentages between 90.04-93.33% (Table 1). This high multimapping percentage can likely be attributed to the Trinity assembly process, which is conservative in its separation and identification of unique transcripts, producing high intra-assembly redundancy (27).

**Table 1:**
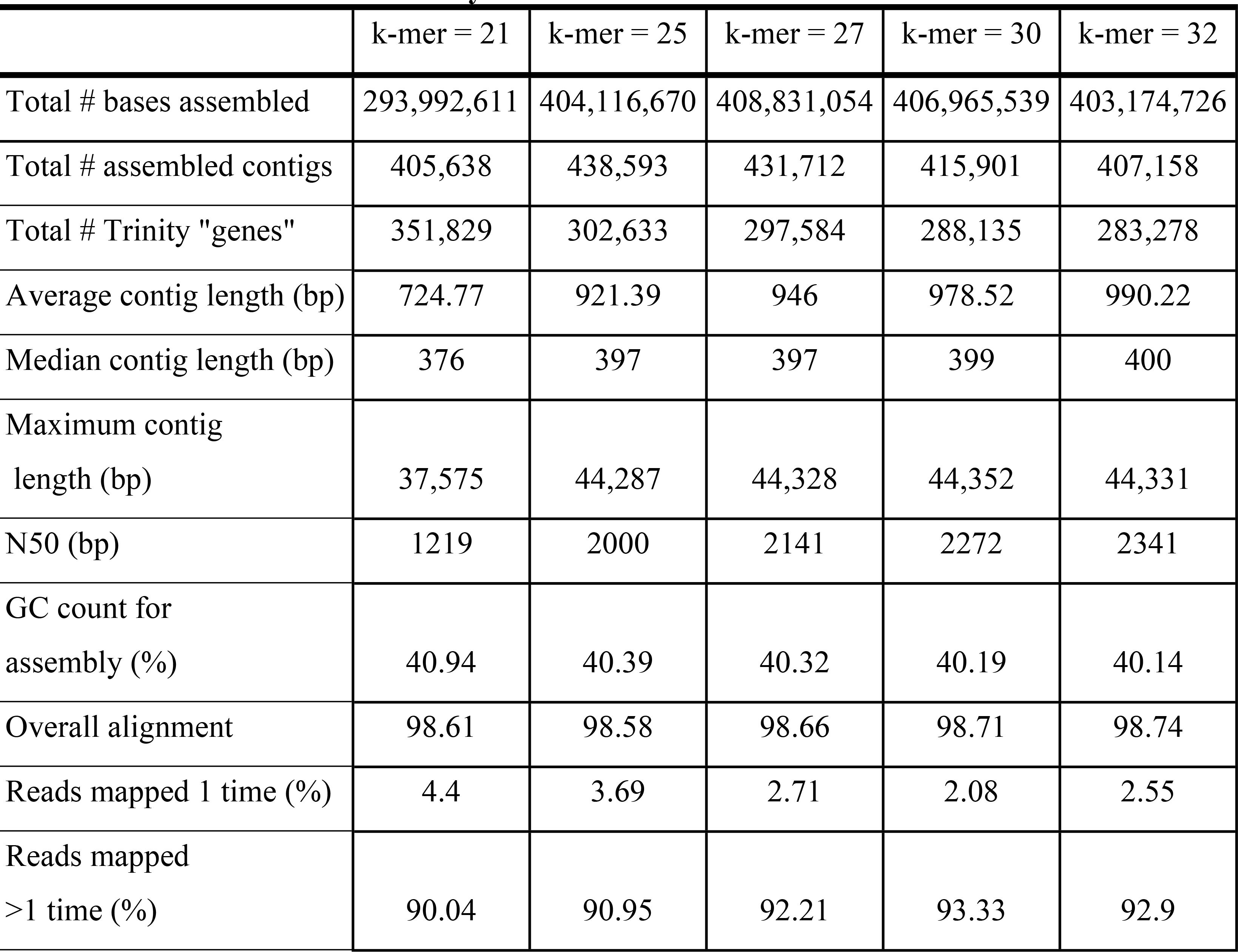
Individual k-mer assembly details. Summary metrics for five different de novo transcriptomes built with five different k-mer lengths.

|  | k-mer = 21 | k-mer = 25 | k-mer = 27 | k-mer = 30 | k-mer = 32 |
| --- | --- | --- | --- | --- | --- |
| Total # bases assembled | 293,992,611 | 404,116,670 | 408,831,054 | 406,965,539 | 403,174,726 |
| Total # assembled contigs | 405,638 | 438,593 | 431,712 | 415,901 | 407,158 |
| Total # Trinity "genes" | 351,829 | 302,633 | 297,584 | 288,135 | 283,278 |
| Average contig length (bp) | 724.77 | 921.39 | 946 | 978.52 | 990.22 |
| Median contig length (bp) | 376 | 397 | 397 | 399 | 400 |
| Maximum contig length (bp) | 37,575 | 44,287 | 44,328 | 44,352 | 44,331 |
| N50 (bp) | 1219 | 2000 | 2141 | 2272 | 2341 |
| GC count for assembly (%) | 40.94 | 40.39 | 40.32 | 40.19 | 40.14 |
| Overall alignment | 98.61 | 98.58 | 98.66 | 98.71 | 98.74 |
| Reads mapped 1 time (%) | 4.4 | 3.69 | 2.71 | 2.08 | 2.55 |
| Reads mapped >1 time (%) | 90.04 | 90.95 | 92.21 | 93.33 | 92.9 |

The five transcriptomes were combined to generate a transcriptome with a total of 2,099,002 contigs (Figure 1), presumably many of which were redundant. We used the EvidentialGene *tr2aacdsmRNA* classifier to filter the redundancies within our transcriptome, which were present due to both intra- and inter-assembly redundancies (21). The EvidentialGene program employs an algorithm that operates on the open reading frames of the contigs to generate a non-redundant transcriptome containing the optimal set of transcripts based on biological relevance and coding potential (21, 28). This program is often used in ‘over-assembly’ procedures where multiple assemblies are combined (29–31). With our multi-k-mer assembly, EvidentialGene produced a main ‘okay’ set, containing 55,895 contigs, and an alternative ‘okalt’ set, containing 143,364 contigs, which were combined to produce a final transcriptome with a total of 199,357 contigs, reducing the overall number of contigs by 90.5%. BLAST searches across all the contigs yielded matches for 127,324 transcripts, a 63.87% BLAST hit rate for the entire transcriptome. The number of Trinity predicted genes after running EvidentialGene dropped slightly to 132,972. The multimapping percentage was reduced from approximately 90% to around 21%.

**Figure 1.**
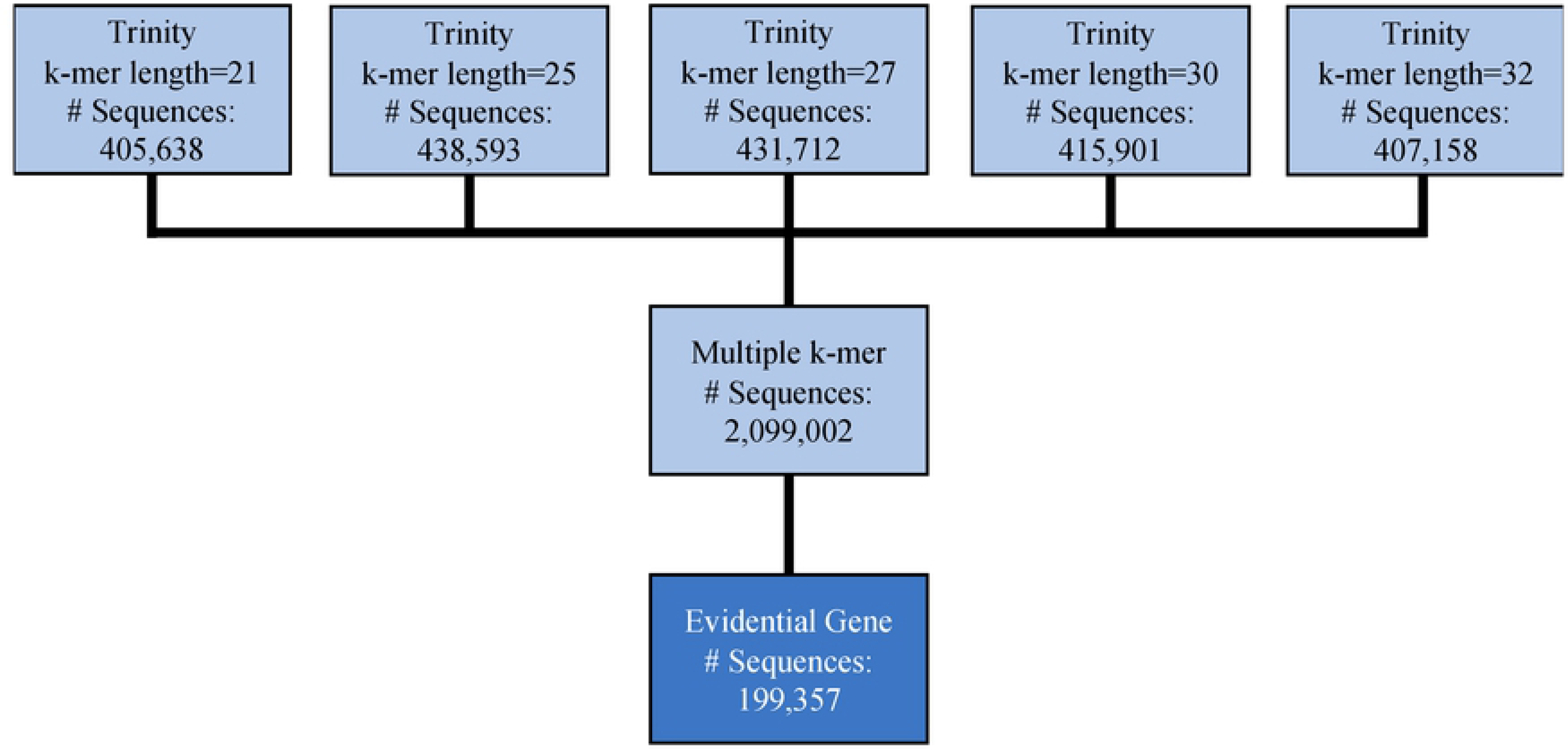
Summary of workflow for multi k-mer assembly.

To check the accuracy of the sequences predicted in the transcriptome, we used Sanger sequencing to independently confirm the sequences of six randomly selected transcriptome transcripts. Of these six, four of them were predicted to contain complete open reading frames (ORFs), and two were missing the 3’ end. We analyzed 14,299 nucleotides of 15,478 predicted base pairs (92%). The number of substitutions (16), insertions (84), and deletions (0) were noted; overall, these differences accounted for approximately ∼0.1% of the sequenced nucleotides (data not shown). A few additional randomly selected sequences were highly repetitive and were not amenable to Sanger sequencing; we did not proceed with an analysis of any of these candidates. **Differential expression during compensatory plasticity**

To determine genes that were differentially regulated during compensatory plasticity, the reads for each of the 16 Illumina libraries, which excluded the two outliers and three backfill libraries (see Methods), were mapped back to our multiple k-mer transcriptome creating a counts matrix. Pairwise comparisons of normalized counts data from deafferented *vs*. control crickets were performed at each time point using both algorithms, EdgeR and DESeq2 (See Supplemental Materials). The distribution of differentially expressed genes was initially visualized using volcano plots (Figure 2). These plots revealed slightly different distributions of upregulated versus downregulated genes between the two programs. Overall, however, we saw strong correlations between these two programs for all time points (Figure 3), with the exception of a few of the high fold-change candidates. Within this range, EdgeR was consistently more conservative than DESeq2, which was especially true for a small number of upregulated candidates (Figure 3a-c).

**Figure 2.**
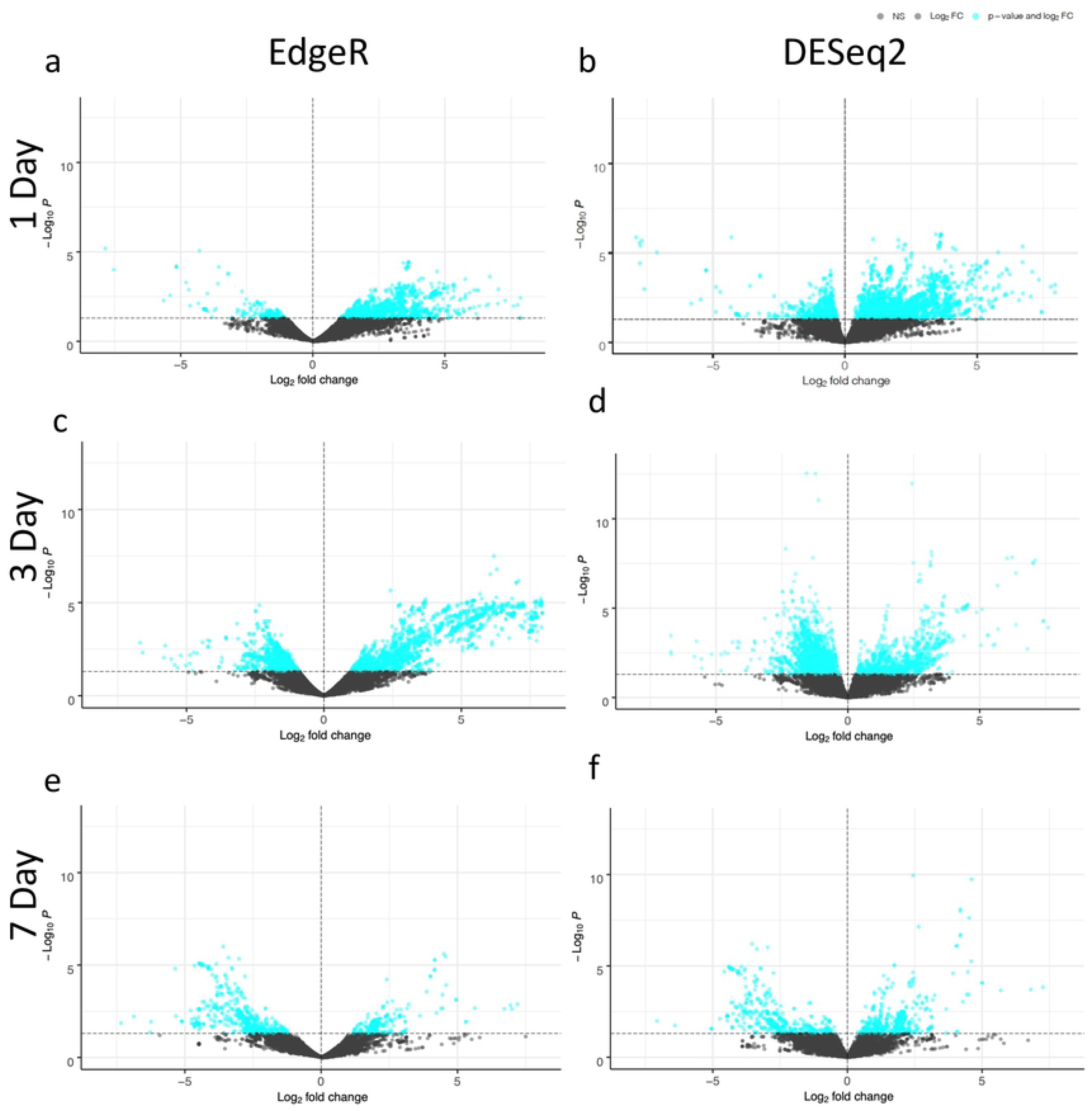
Volcano plots of differential gene expression in *G. bimaculatus* prothoracic ganglia at one day (a,b), 3 days (c,d) and 7 days (e,f) after deafferentation. The horizontal dotted line marks a p-value of 0.05, and the vertical dotted line marks no predicted fold change. Each point represents a contig determined to be differentially regulated by EdgeR (a,c,e) or DESeq2 (b,d,f). Blue points represent the contigs determined to be significantly regulated.

**Figure 3:**
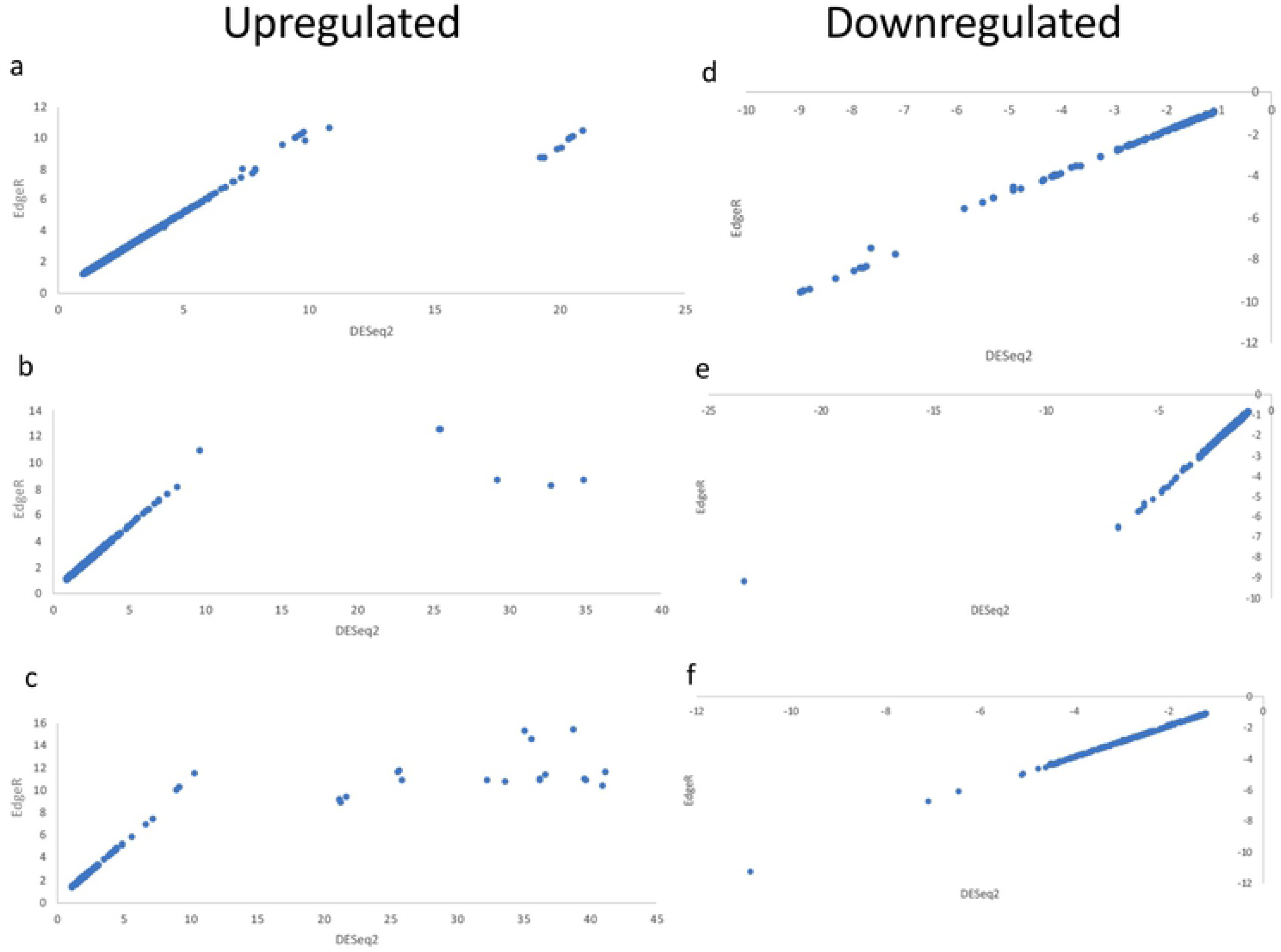
Correlation of fold-changes predicted by EdgeR and DESeq2 for upregulated transcripts (a,b,c) and downregulated transcripts (d,e,f) at one(a,d), three (b,e), and seven (c,f) days.

The majority of the transcripts were upregulated in the 2 to 4-fold range at one day (59% of the transcripts), three days (55% of the transcripts), and seven days (45% of the transcripts). The next largest group of transcripts was upregulated at 0 to 2-fold at one day (33% of transcripts), three days (41% of transcripts), and seven days (39% of transcripts). For those candidates that were downregulated, a majority of them at one day (63%) and three days (86%) were downregulated less than 2-fold. At seven days, the bulk of candidates (70%) were downregulated 2 to 4-fold. Analysis of 10 of the transcripts at each time point with the largest fold changes revealed that most were unidentified and did not match anything in the NCBI database when BLASTed. A few of these transcripts did have BLAST hits, such as a mucin- 5AC-like (down at one day), larval cuticle protein-3-like (down at seven days), and hypothetical accessory gland protein (up at three and seven days).

Using EdgeR, 261 genes were found to be downregulated at one day post-deafferentation, 1,675 genes were downregulated at three days post-deafferentation, and 580 genes were downregulated at seven days post-deafferentation (Figure 4a). Additionally, 2,234 genes were determined to be upregulated at one day post-deafferentation, 1,860 genes upregulated at three days post-deafferentation, and 290 genes upregulated at seven days post-deafferentation (Figure 4b).

**Figure 4.**
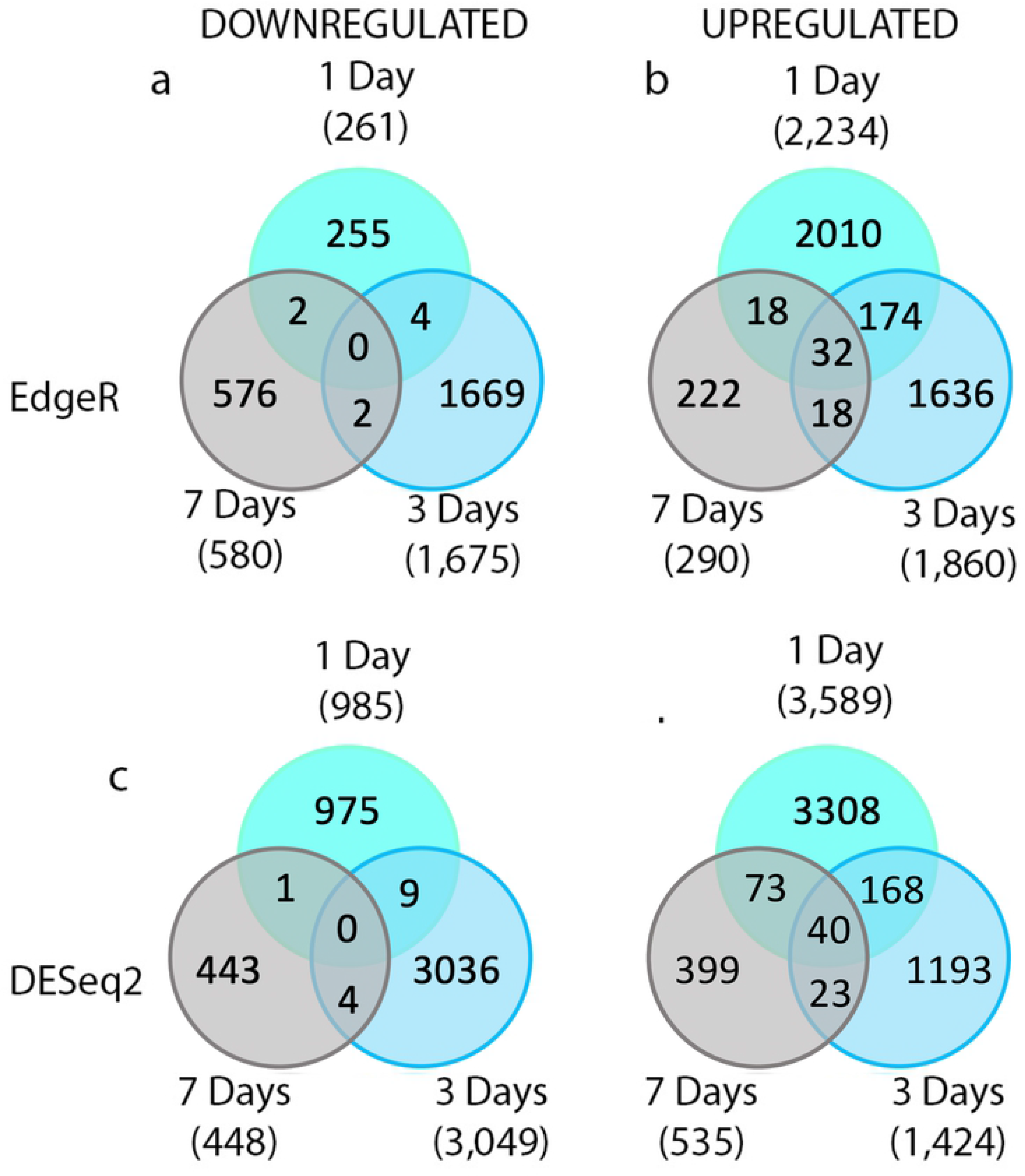
Differentially regulated genes across three time points. Similar patterns in relative numbers of differentially regulated genes are observed between the two programs. a) EdgeR identified downregulated genes b) EdgeR identified upregulated genes c) DESeq2 identified downregulated genes d) DESeq2 identified upregulated genes.

A similar pairwise comparison of deafferented versus control crickets was performed using the DESeq2 software and revealed that 985 genes were downregulated at one day post-deafferentation, 3,049 genes were downregulated at three days, and 448 genes were downregulated at seven days (Figure 4c). Additionally, 3,589 genes were upregulated at one day post-deafferentation, 1,424 genes were upregulated at three days, and 535 genes were upregulated at seven days (Figure 4d).

From these sets, simple comparisons were created to determine the number of genes upregulated and downregulated across multiple timepoints. In EdgeR, there were four genes downregulated at one and three days, two genes at one and seven days, two genes at three and seven days, and 0 genes differentially downregulated across all three time points (Figure 4a). For the upregulated genes, there were 174 genes differentially regulated at one and three days, 18 genes at one and seven days, 18 genes at three and seven days, and 32 genes at all three time points (Figure 4b). Comparing the DESeq2 sets of genes across multiple timepoints showed that there were nine genes downregulated at one and three days, one gene downregulated at one and seven days, four genes downregulated at three and seven days, and 0 genes downregulated at all three timepoints (Figure 4c). Additionally, 168 genes were found to be upregulated at one and three days, 73 genes at one and seven days, 23 genes at three and seven days, and 40 genes at all three time points (Figure 4d).

Finally, simple comparisons were performed between EdgeR and DESeq2 with genes determined to be differentially expressed at each of the three times points. For downregulated genes, 180 were identified at one-day post deafferentation, 1,604 at three days, and 367 at seven days. For upregulated genes, 2,099 were identified at one-day post deafferentation, 1,043 at three days, and 272 at seven days (Figure 5). The genes found to be differentially regulated by both programs were used for further analysis.

**Figure 5.**
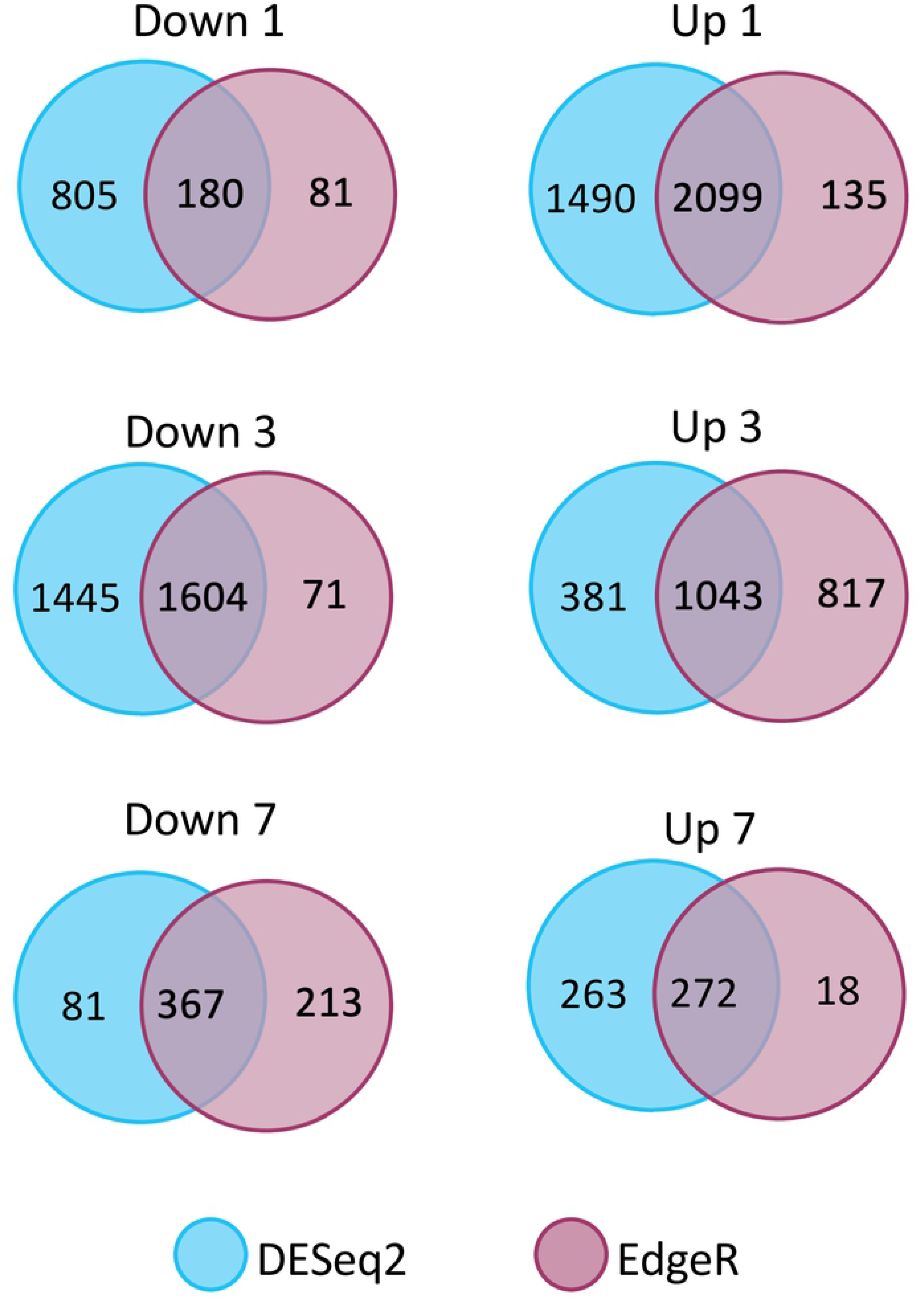
Differentially regulated genes compared across the two analytical programs, DESeq2 and EdgeR. The number of genes found to be differentially regulated by both programs varies by condition.

DESeq2 and EdgeR are the leading programs used for the analysis of RNA-Seq data, with thousands of reports both using these methods for analyzing differential expression and comparing their computational methods (32, 33). While both operate under the hypothesis that the majority of genes are not differentially expressed, they employ different computational methods, especially with respect to the normalization process, to determine differentially expressed genes (33). EdgeR and DESeq2 both use a normalization by distribution method, but EdgeR relies on the Trimmed Mean of the M-values method, whereas DESeq2 uses a Relative Log Expression method (34–37). Since different methods rely on differing assumptions in order to identify differentially expressed genes, the results will vary slightly. One experiment comparing EdgeR and DESeq2 found relatively similar lists of differentially expressed genes produced by the two programs, with EdgeR producing more conservative, smaller gene lists (32). In this study we decided to use two different programs to conduct the differential expression analysis in order to create a smaller, more conservative set of genes for future functional analyses. Out of the six comparisons between EdgeR and DESeq2 (upregulated and downregulated at one, three, and seven days post injury), four out of the six resulted in EdgeR producing a smaller set of genes than DESeq2 (Figure 5), in line with the study by Raplee and colleagues (32). Although the two programs generated varying numbers of differentially regulated genes, similar patterns in relative numbers were observed. Both programs showed a decrease in the number of genes upregulated across time. For the downregulated genes, a peak in the number of differentially regulated genes was found at three days post injury. This similarity was expected given the relative similarity and previous studies of both analysis programs.

### BLAST and Gene Ontology Annotations

Once we had identified a conservative set of transcripts predicted to be differentially regulated, we used BLAST2GO (38) to try to identify them. Not all the transcripts inputted into the BLAST2GO program resulted in BLAST hits and/or GO annotations. At one day downregulated, 71% of genes had both BLAST and GO results and an additional 10% had only BLAST hits. At three days downregulated, 36% of genes had both BLAST and GO results and an additional 6% had only BLAST hits. At seven days downregulated, 31% of genes had both BLAST and GO results and an additional 6% had only BLAST hits. For the one day upregulated, 53% of genes had both BLAST and GO results and an additional 10% had only BLAST hits. At three days upregulated, 59% of genes had both BLAST and GO results and an additional 15% had only BLAST hits. At seven days upregulated, 50% of genes had both BLAST and GO results and an additional 15% had only BLAST hits (Figure 6).

**Figure 6.**
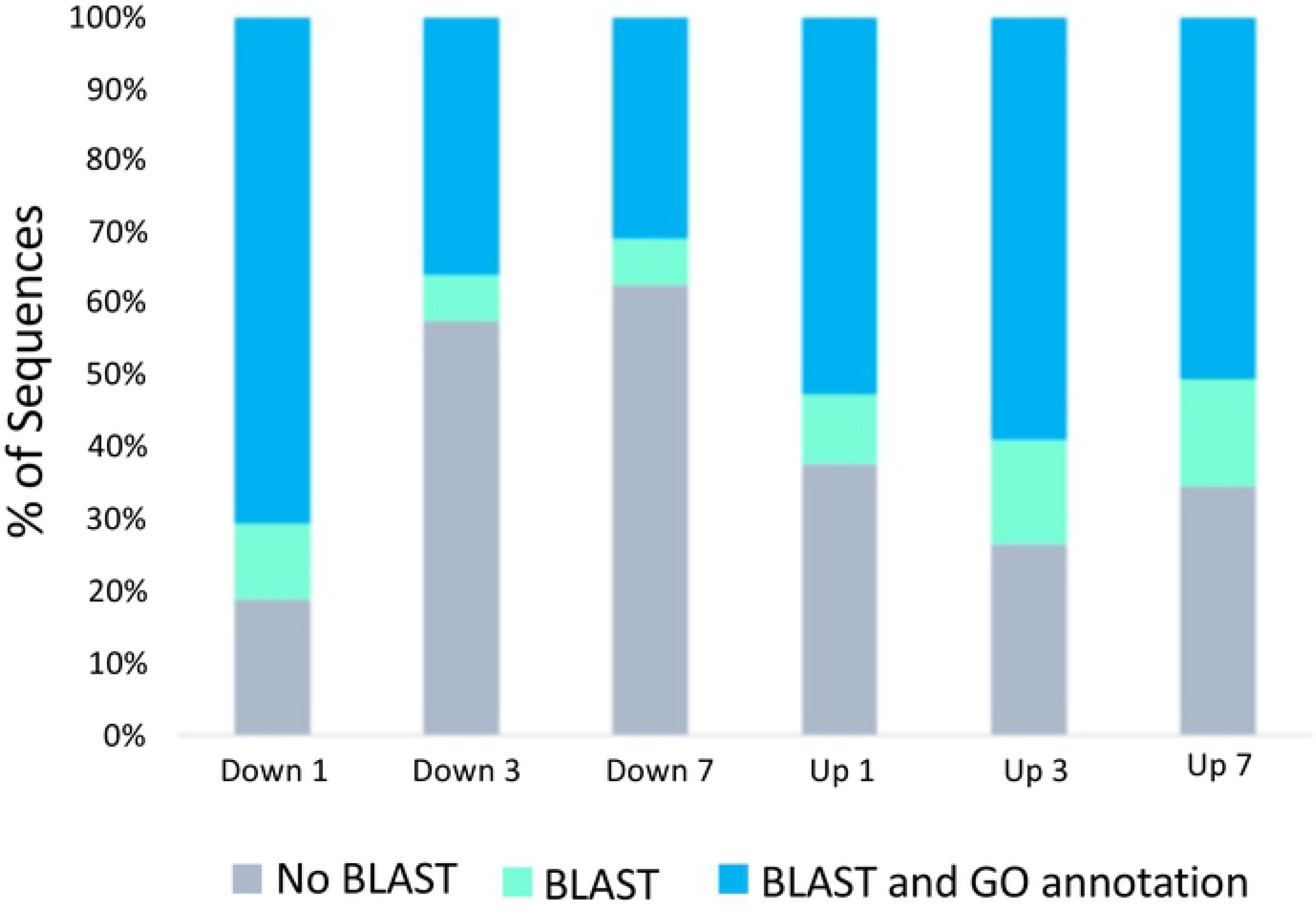
Percentage of sequences with no BLAST hits, BLAST hits, and BLAST hits with additional GO term mapping. Distribution of sequences varies across times points and regulation.

For the six lists of differentially expressed transcripts, there was a range between 37-81% of transcripts having BLAST hits against the nr database and 23-62% against the manually curated and annotated Swiss-Prot database. After mapping with GO terms, this was reduced to about 31-59%. This left close to half of the differentially regulated transcripts with no functional information. These transcripts could represent uncharacterized proteins, which may or may not be playing an important role in the compensatory growth response. Since we performed a BLASTx looking at proteins, it is also possible that these transcripts are non-coding RNAs. Although polyA selection was used as part of the RNA-Seq process, this may not be completely efficient in removing all non-coding RNAs, specifically long non-coding RNAs (39, 40). Finally, it is also possible that there were issues within these transcripts themselves, either due to an error during the assembly process or the sequences being too short to be matched with confidence. Regardless, we completed no further analysis of these transcripts.

One set of intriguing proteins that were found to be upregulated at one day were members of the Protein Yellow family. Our transcriptome predicted the upregulation of 15 different Protein Yellow transcripts that appear to consist of at least eight different splice variants of Protein Yellow-f (data not shown). We used qPCR to confirm the upregulation of these transcripts in independent samples and found weak support for this upregulation (p=0.1; data not shown). The primers used were designed to target all eight splice variants identified in our transcriptome, but because all the splice variants of Protein Yellow-f in the cricket are not well-characterized, we may have inadvertently captured additional splice variants in this analysis that are not differentially expressed. Future experiments confirming the enrichment of each of these splice variants along with functional validation will be necessary.

Protein Yellow belongs to the Major Royal Jelly protein family and are secreted proteins found in the extracellular region (41). Protein Yellow was first characterized in *Drosophila melanogaster* for its role in pigmentation (42). Other research in honeybees revealed the importance of Royal Jelly proteins in development and social behavior in addition to a possible role in the CNS (41, 43). However, the function of these Yellow/Royal Jelly proteins is not completely understood (42). While the role of these proteins in crickets is unclear, they were statistically differentially regulated and would be an interesting molecular family to investigate for their role in deafferentation-induced plasticity.

### GO Term Distributions

Based on a preliminary grouping of GO terms by the three root classes, it appeared that several classes of GO terms were found to be associated with our differentially expressed genes (Figure 7). While the top five represented GO terms encompassed most of the GO terms in the cellular component category, there was a much broader range of GO terms represented in the molecular function and biological process categories

**Figure 7.**
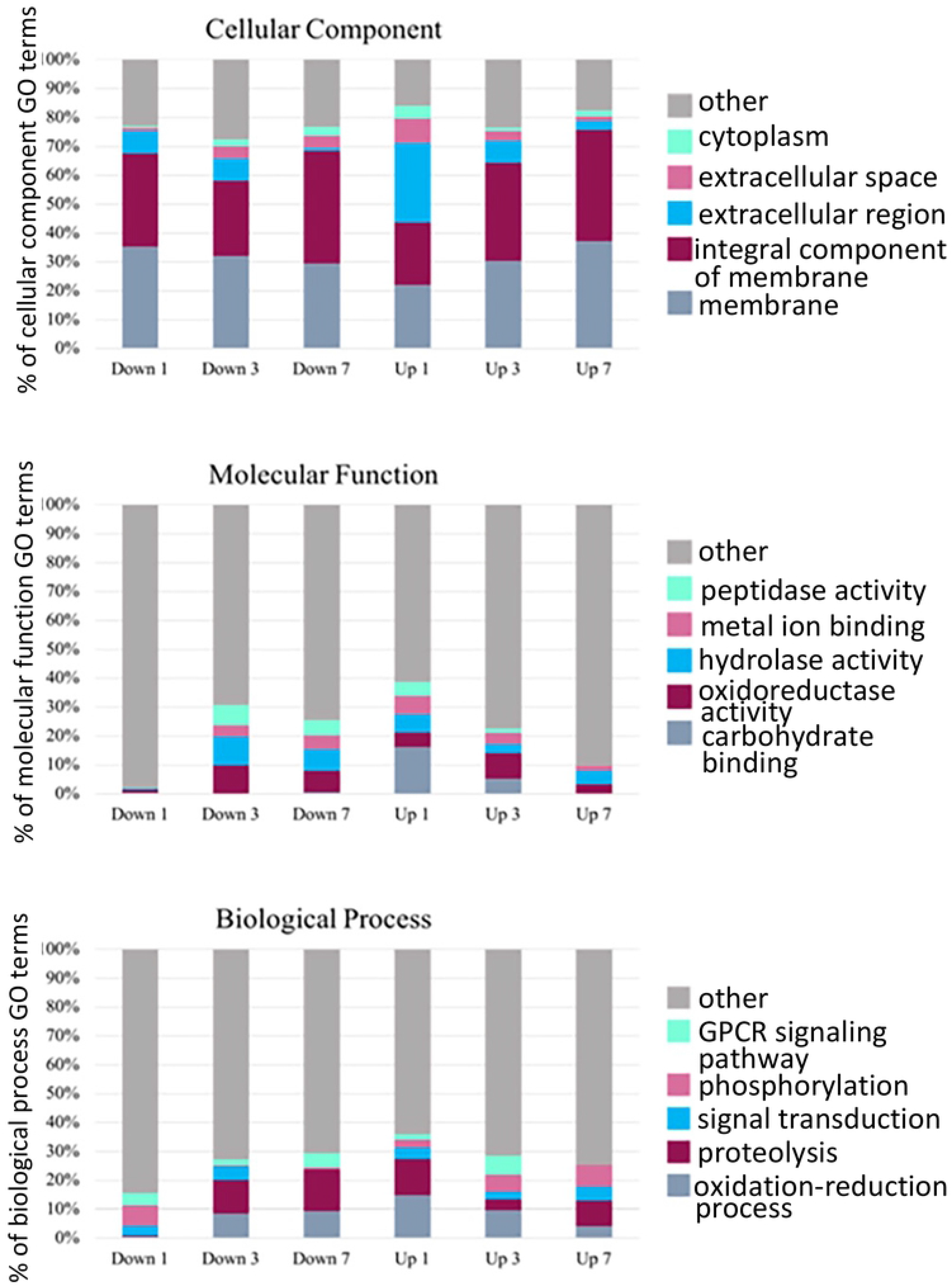
GO term analysis organized into the three root classes: cellular component (CC), molecular function (MF), and biological process (BP). The top 5 represented GO terms across all time points in each class are represented. Many highly represented GO terms were found in the CC class whereas a broader range of GO terms were found in the MF and BP classes.

Web Gene Ontology Annotation Plot (44, 45) was used to plot a broader distribution of GO terms and visually compare annotations among timepoints (Figure 8). Cellular component, molecular function, and biological process were displayed on traditional WEGO histograms (Figure 8a, b). The percentage of genes indicates the percentage of the genes within a given list that were annotated with the given GO term or one of the child nodes of the term. GO terms with higher percentage representation included terms describing membrane-related components as well as terms related to catalytic activity, binding, and metabolic and cellular processes.

**Figure 8.**
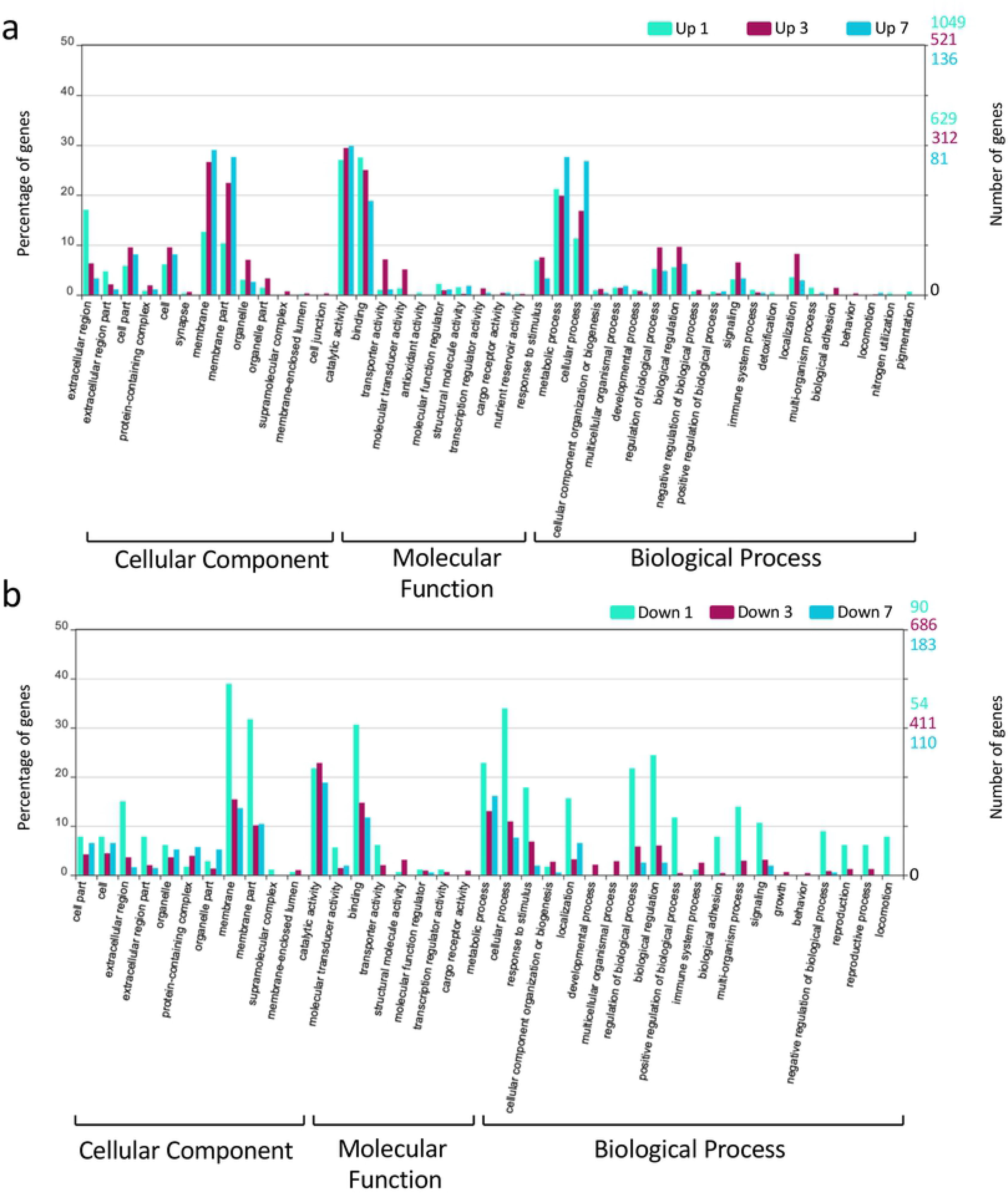
WEGO histograms with the distribution of Gene Ontology terms grouped by cellular component, molecular function, or biological process. a) GO terms associated with upregulated genes across all time points. b) GO terms associated with downregulated genes across all time points. Percentages are noted on the left and the number of genes within the given list that were annotated with the GO term/child term are noted on the right. On the right axis, the top numbers (turquoise) correspond to the one-day data, the middle numbers, (magenta) corresponds to the three-day data, and the bottom numbers correspond to the seven-day data (teal).

### Gene Ontology Categories

We examined whether any of the candidate GO terms we identified here were associated with injury-related plasticity paradigms identified in other species. For example, perhaps successful regeneration after injury depends on the recapitulation of developmental proteins that promote neurogenesis (46) or guide axons and dendrites (18). If these molecular strategies were important for the plasticity observed in the cricket, we would predict that we might see changes in the expression of transcription factors involved in neurogenesis and/or the regulation of several classes of guidance cues normally involved in midline control in insect embryos. When searching our differentially regulated candidates, a few genes downregulated at three days were associated with GO terms that were related to neurogenesis (**GO:0007465**: R7 cell fate commitment and **GO:0045466**: R7 cell differentiation). We found only one transcript that was annotated with an axon guidance-related GO term, which was identified as a “twitchin-like protein,” (Table 2). Twitchin/Unc-22 is a large protein kinase thought to be important in muscle development and function (47).

**Table 2:**
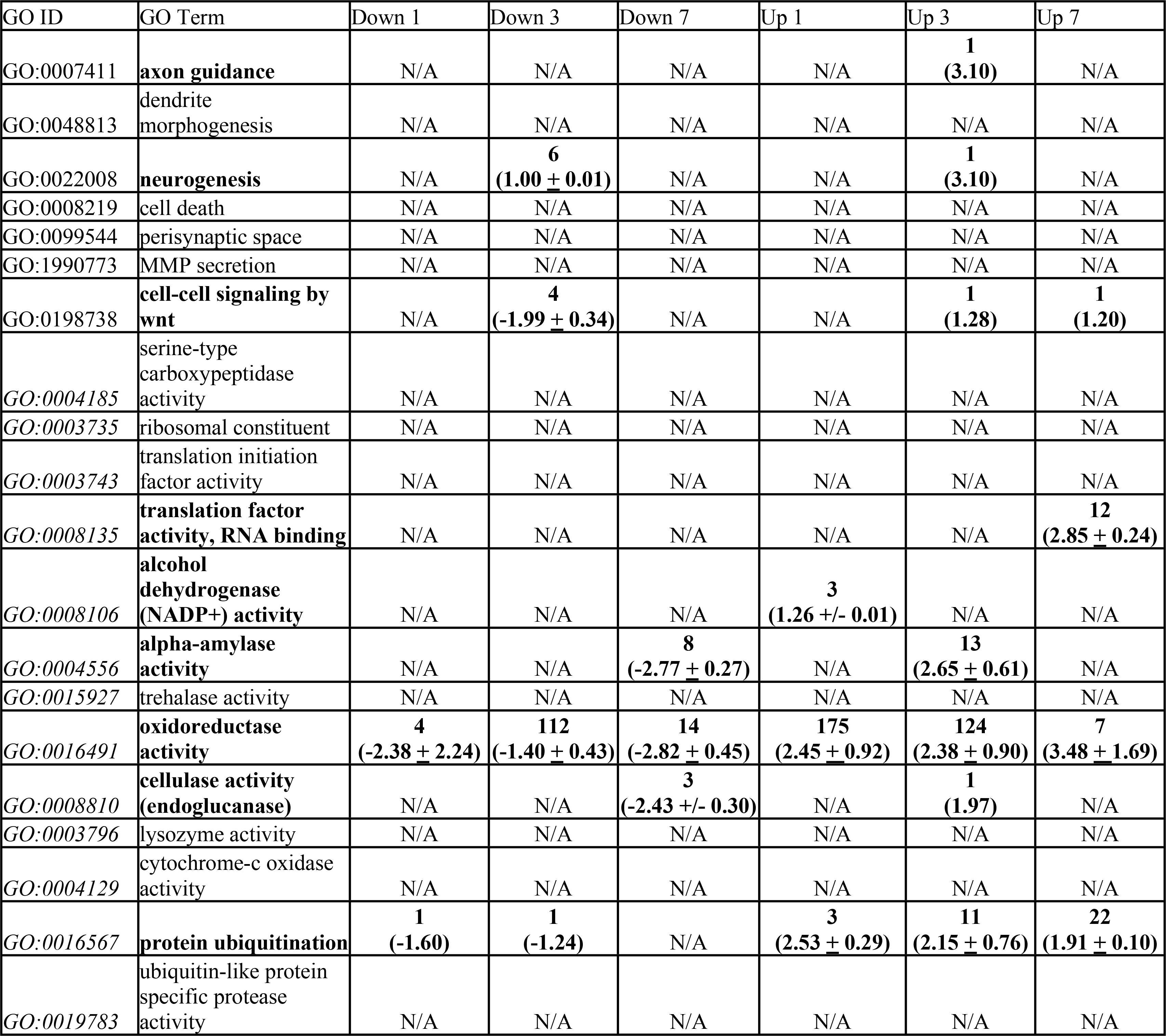
Evaluation of the number and fold-change of GO terms of interest. Number of transcripts associated with the selected GO terms, including any child term of the GO term, at each time point. For each represented GO term, the average +/- standard deviation of the log2foldchange across the transcripts is given in parentheses. GO Terms in bold indicate significant changes in expression. GO IDs in italics were selected because similar transcripts were present in a prior suppression subtractive hybridization study (54).

An initial study of our original single k-mer transcriptome explored this developmental recapitulation hypothesis by mining the transcriptome for the presence of various guidance molecules (15). Though transcripts corresponding to the signaling families, slit, netrin, ephrin, and semaphorin were identified within this adult transcriptome, the BLAST and Gene Ontology analyses performed here did not identify an abundant number of guidance molecules as differentially regulated. Despite this result, however, it is important to note that the transcriptome and differential expression analysis were performed on the whole prothoracic ganglion, which could mask important changes that occur in single cells after deafferentation, such as in ascending neuron 1 and 2 (AN-1 and AN-2). Single cell RNA-Seq analysis of the ANs could help to determine whether there are changes in expression occurring on a smaller anatomical level.

Based on results from different types of injury model systems in other organisms, additional functional categories that we hypothesized could change during the compensatory growth response were those related to apoptosis (48, 49) and Wnt signaling (50). In our differentially expressed gene sets we did not find enrichment in terms related to apoptosis. Searching our results for Wnt-related GO terms, revealed a few genes annotated with Wnt pathway members at 3 days post deafferentation (Table 2). These genes had a top BLAST hit of atrial natriuretic peptide-converting enzyme isoform X1, Frizzled-4, and secreted frizzled-related protein 5-like.

We looked for the presence of a number of additional groups of proteins that influence neuronal morphogenesis, plasticity, or remodeling (Table 2). For example, the matrix metalloproteases (MMPs) are required for axonal guidance (51) as well as dendritic remodeling during metamorphosis in *Drosophila melanogaster* (52). Importantly, the expression of some MMPs appear to contribute to poor recovery after spinal cord injury in mammals (53). We did not find enrichment in any of these terms at any time point (Table 2), indicating that the injury-induced anatomical plasticity in the cricket may rely on different pathways than have been identified in other species. Furthermore, it is notable that factors, such as MMPs, that can restrict growth or contribute to pruning in other organisms are not upregulated upon injury in the cricket.

Several GO terms associated with the candidates found in a previous subtraction hybridization study (54) were also found to be differentially expressed in the present experiment, often showing significant changes in expression at the three- and seven-day post-deafferentation time point (Table 2, bold). These include oxidoreductase, alpha-amylase, endoglucanase, and alcohol dehydrogenase. As noted by Horch and colleagues (54), many of these enzymes have been observed in the hemolymph of insects and play a role in metabolism and immune response. Although it is possible that these findings are due to contaminants during the extraction of the prothoracic ganglion, the results would imply that the extractions differed between control and experimental animals in multiple experiments. Given that the differential expression of several enzyme transcripts was found both in this study and in our former suppression subtractive hybridizaiton (SSH; Ref. 54) study, it is less likely that these enzymes are artifact or contamination effects. Particularly, several differentially regulated transcripts were associated with oxidoreductase activity across all time points. The BLAST hits of these transcripts showed some of the enzymes to be retinal dehydrogenases. Retinal dehydrogenase along with alcohol dehydrogenase, another regulated GO term, are involved with the production of retinoic acid. Retinoic acid has been shown to be involved with development, regeneration, synaptic plasticity, and neurite outgrowth (55–57) implying that regulation of retinoic acid production may influence these processes. Another class of oxidoreductases that appeared abundant within the BLAST hits was the cytochrome P450 family. Cytochrome P450 is a superfamily of monooxygenase enzymes and several families of cytochrome P450 exist in insects. These enzymes are known to have a variety of functional roles in insects including growth and development (58). Cytochrome P450 has also been shown to regulate ecdysone signaling in insects, including crickets (59, 60). Ecdysone signaling is crucial for developmental processes and morphogenesis, but has also been shown to be important in the dendritic remodeling of *Drosophila melanogaster* sensory neurons (52, 59). While these protein families represent some of the transcripts annotated with “oxidoreductase activity”, given the wide range of such transcripts, it is difficult to discern the role of all of the regulated proteins.

### Gene Ontology Enrichment Analysis

Metascape (61) was used to determine enriched GO terms across the differentially expressed gene lists. Differentially expressed genes were first reBLASTed against the curated Swiss-Prot database to retrieve appropriate gene identifiers. Similar ratios of BLAST hit percentages across timepoints were observed against Swiss-Prot as with the nr database, however, the percentage of genes with BLAST hits was lower when BLASTed against Swiss- Prot versus the nr database (Table 3).

**Table 3:** Comparison of the percentage of genes with BLAST hits in the nr database vs. Swiss-Prot.

|  | nr | Swiss-Prot |
| --- | --- | --- |
| Down 1 | 81% | 59% |
| Down 3 | 42% | 36% |
| Down 7 | 37% | 23% |
| Up 1 | 62% | 59% |
| Up 3 | 73% | 62% |
| Up 7 | 65% | 57% |

Enrichment analysis by Metascape showed the most enriched terms at three days post-deafferentation across both upregulated and downregulated genes. Examining the multi-list analysis, there were 22 enriched GO terms found including those related to morphogenesis, extracellular space, and neuron fate commitment (Figure 9). No enriched GO terms were found in the upregulated gene set at seven days post-deafferentation.

**Figure 9.**
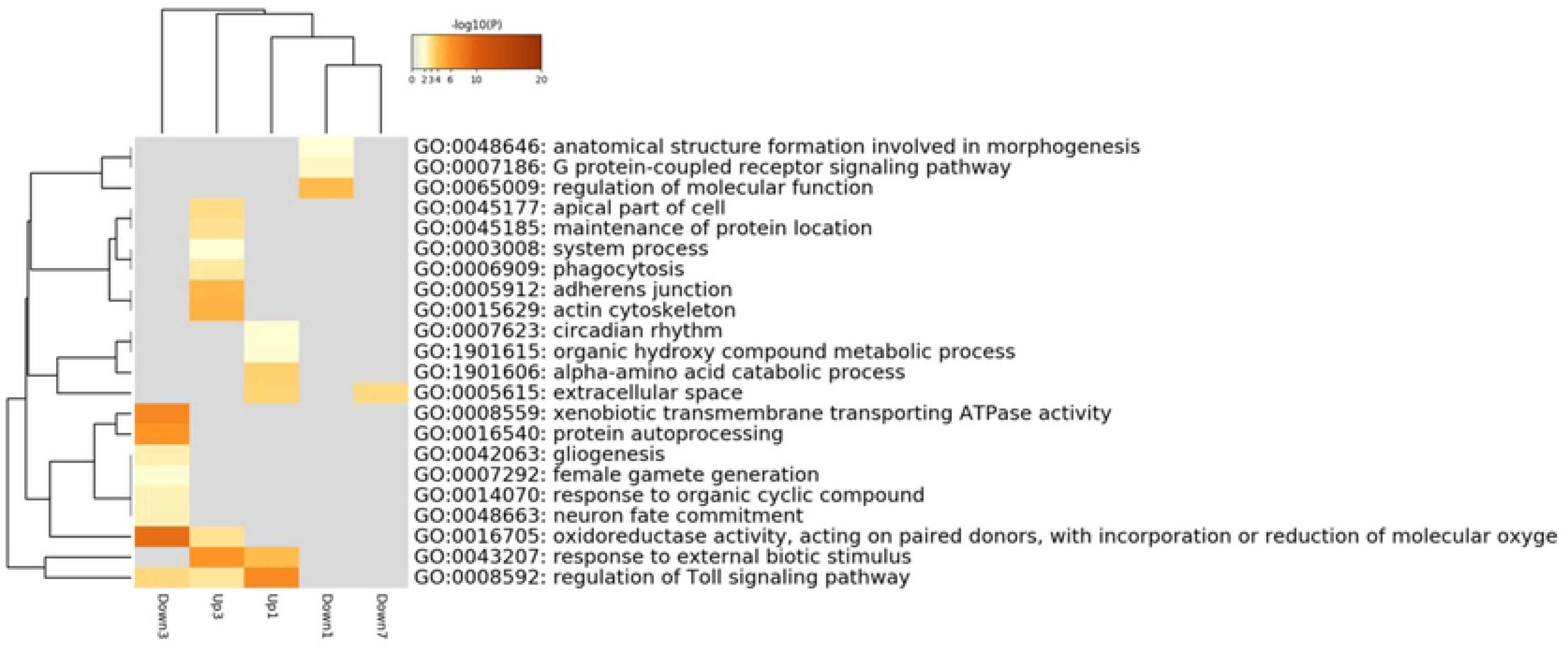
Heatmap of enrichment terms as determined by Metascape. Colored by p-value as indicated at the top.

One category of interest that was revealed in this enrichment analysis was the GO Term “Regulation of Toll-signaling pathway.” Our results indicate over 40 different Toll-signaling-related transcripts were differentially expressed, including Toll receptors and the serine proteases, Snake and Spirit. The differential regulation of a relatively large number of transcripts related to Toll-signaling encourages us to generate future hypotheses focused on the role of this pathway in the injury-induced plasticity of the auditory system. At that point, validation of sequence, function, and expression levels will be necessary, especially because several of the identified candidates are likely splice variants of individual genes.

Toll receptors are most commonly associated with their function in immunity and development, however, research in *Drosophila melanogaster* suggests that they may also play a role in regulating cell number, connectivity, and synaptogenesis (62). Activation of Toll receptors can regulate cell number through either neuroprotective or pro-apoptotic functions, depending on the receptor type. These functions of Toll receptors were shown to exist in both development and adulthood (63). Toll receptors, specifically Toll-6 and Toll-7, have also been shown to have neurotrophic receptor-like functions through their ability to bind multiple ligands, including neurotrophin-like proteins in invertebrates (64). Neurotrophins are known to regulate cell proliferation and neuronal survival and development, thereby suggesting an important role for Toll receptors in neuronal systems (63, 64). Furthermore, in *Drosophila melanogaster* the receptor Toll-8 was shown to positively regulate synaptic growth through a retrograde neurotrophin-like signaling mechanism (65). These studies show that Toll signaling may play an important role in regulating structural plasticity in invertebrates and, given their enrichment in our differentially regulated gene set, may be crucial to the dendritic reorganization observed in the cricket.

### Conclusions

Unilateral tympanal organ removal in the cricket, *Gryllus bimaculatus*, leads to a robust reorganization of dendrites in the auditory system of the prothoracic ganglion. This novel growth and *de novo* synapse formation restores the ability of the deafferented neurons to respond to sound. Our transcriptomic analyses identified thousands of transcripts up- and down-regulated after deafferentation. We highlight transcriptional changes related to protein translation and degradation, enzymatic activity, and Toll signaling that appear to be enriched after deafferentation. The data presented here allows the development of targeted hypotheses that could elucidate the mechanisms responsible for the deafferentation-induced synaptic plasticity in the auditory system of crickets. The mechanisms at play here can be compared and contrasted with those identified in the terminal ganglion of the cricket after unilateral loss of a cercus (66).

## Methods

### Animals, injury, and library preparation

Prothoracic ganglia from approximately 60 adult, male Mediterranean field crickets, *Gryllus bimaculatus* were harvested and 21 individual ganglia were ultimately used as the sources of RNA for this transcriptome (15). Male crickets that were adults for 3-5 days received either a control amputation of the distal segment of the left tarsus (“foot chop” control crickets), or the left prothoracic leg was severed mid-femur removing the auditory organ and deafferenting the ipsilateral central auditory neurons (“deafferented” experimental crickets). Males were chosen due to the potential sexual dimorphism in rates of dendritic growth after deafferentation (22). Several crickets were prepared for backfill as previously described (15). Prothoracic ganglia were removed from crickets 1, 3, or 7 days after amputation at the femur or tarsus, or 18 hours post-backfill, and total RNA was purified as previously described (Figure 10).

**Figure 10.**
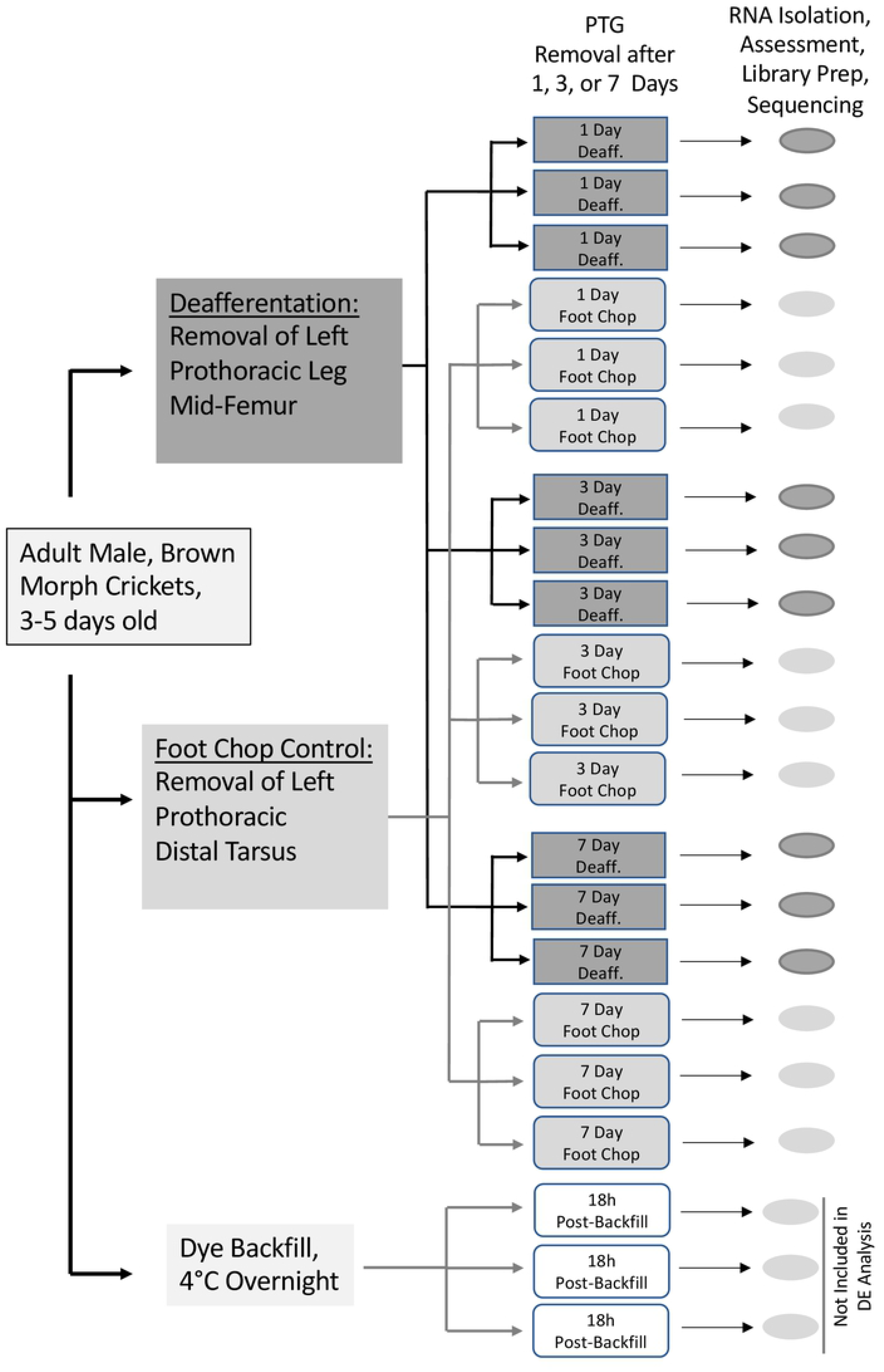
Summary of experimental design. 21 PTG ganglia were removed from “deafferented” or “foot chop control” animals 1, 3, or 7 days-post injury. Three additional animals were backfilled 18 hrs prior to PTG removal. RNASeq data from these animals were included in the assembly but not in the differential expression.

The QIAGEN RNeasy Lipid Tissue Minikit was used to purify total RNA from each sample individually. RNA concentrations were assessed after TURBO DNA-free treatment (Ambion by Life Technologies) with a nanospectrophotometer (Nanodrop, Thermo Fisher Scientific) or a fluorometer (Qubit, Thermo Fisher Scientific). An Agilent 2100 Bioanalyzer (Applied Biosystems, Carlsbad, CA) was used to further assess sample quality. Based on evaluation of RNA quality and concentration of individual ganglion samples, the best 3 samples for each condition were selected for sequencing. Standard Illumina paired-end library protocols were used to prepare samples. The Illumina Hiseq 2500 platform, running v4 chemistry to generate ∼25M paired end reads of 100bp in length for each sample, was used to sequence the RNA (15).

### Transcriptome assembly

Trinity software (Trinity-v2.6.5) was run using previously processed and filtered reads of prothoracic ganglion libraries (15). A multi-k-mer assembly was created by building five *de novo* transcriptomes using a single k-mer length (21, 25, 27, 30, 32) and subsequently combining them. The following parameters were used: minimum contig length of 200, library normalization with maximum read coverage 50, RF strand specific read orientation, maximum memory, 250GB, and 32 CPUs. Individual assemblies were analyzed using the *TrinityStats.pl* script and alignment statistics were obtained using Bowtie2 (v 2.3.4.1). The PRINSEQ interactive program (67) was used to generate additional summary statistics on each assembly.

A k-mer number identity was added to each contig’s Trinity ID, all five assemblies were concatenated, and the Evidential Gene program was used to create a single non-redundant assembly. Evidential Gene relies on the Transdecoder.LongOrfs method, identifies the longest ORFs, removes fragments, and uses a BLAST on self to identify highly similar sequences (98%). The main (okay) and alternative (okalt) sets output from Evidential Gene were combined into a final FASTA file used as the transcriptome for all subsequent analyses. Bam files, sorted bam files, bam index files, and idxstats.txt files were created using samtools (68). This assembly is publicly available at NCBI (Bioproject: PRJNa376023, SUB8325660). The metajinomics python mapping tools (69) were used to generate a counts matrix.

### Coverage Analysis

Samtools was used to extract the sequencing depth at every base position for each contig in every cricket sample, and a python script was used to extract the mean and standard deviation of depth for each contig. The package plotly in R (70) was used to plot the depth of each cricket sample. Graphs were visually compared to determine outliers. Two outliers, 1C1 and 7C2, were removed.

EdgeR and DESeq were used to run the differential expression analysis, (35, 37). The raw read counts generated for each of the libraries, excluding the outliers and the backfill conditions, were used as input to both programs. Similar filtering and normalization functions were used in both programs to exclude any contigs that did not have at least one count per million in at least two libraries. Comparisons between control and deafferented samples were performed at each time point to create lists of upregulated and downregulated genes with a p value cutoff of 0.05. Pairwise comparison results were then compared across time points and were then compared and visualized between the two programs. Another set of lists containing the genes overlapping the two programs was created for continued analysis. The EnhancedVolcano package available in R was used to visualize differential gene expression in volcano plots (71).

### PCR Confirmation

Six sequences were randomly selected for amplification and Sanger sequencing in order to validate the assembly. Sets of primers were designed to obtain the sequence of most of each sequence as predicted by the Trinity assembly. Primers, available on request, were designed for TRINITY21_DN57089_c8_g2_i1.p1 (hypothetical coiled-coil domain protein), TRINITY21_DN58301_c9_g1_i8.p1 (eukaryotic translation initiation factor 4 gamma 3-like), TRINITY25_DN131062_c0_g1_i1.p1 (protein unc-13 homolog 4B), TRINITY27_DN140563_c0_g1_i5.p1 (syntaxin-binding protein 5), TRINITY32_DN141398_c1_g1_i7.p1 (cytochrome P450 301a1), TRINITY21_DN54942_c12_g1_i5.p1 (kinesin light chain). cDNA derived from RNA purified form independent control and deafferented prothoracic ganglion samples was used for PCR. PCR amplicons were gel purified and sequenced, and sequences were aligned and analyzed in Geneious Prime Software (Version 2019.2.3).

### qPCR Validation of Protein Yellow

Quantitative PCR was used to validate predicted expression changes in Protein Yellow candidates. RNA was extracted as described above and reverse transcribed with oligo-dT primers and Superscript III. qPCR primers against Protein Yellow-f (Left: GCGTCTGGCAGAACAGCTCC and Right: CGTGGATGAAGGAGGCGGTG) were designed using a modified version of Primer 3 (version 2.3.7) within Geneious and were chosen so that all potential splice variants would be quantified. Reactions were run on an a QuantSudio3 (ThermoFisher) with Power SYBR Green Master Mix (Thermofisher) following manufacturer’s protocols and with 1M betaine in the master mix due to high GC content (72%) of the amplicon. Annealing temperatures of primers were validated by completing a qPCR using a temperature gradient. PCR efficiencies were checked by running 6, 2-fold serial dilutions of cDNA template, with resulting slope value of -3.213 indicating acceptable efficiency. 2 samples per condition were run in triplicate. Expression values were normalized relative to 2 reference genes (Arm and EF1a), which were identified using RefFinder (72) as the most stable among 6 different reference gene candidates. PCR Miner (73) was used to assess differential expression.

### BLAST Searches

A Perl script was used to extract differentially expressed sequences. The NCBI BLASTx local tool (74) was used to identify proteins similar to the translated nucleotide query sequences. An E-value cutoff of 1e-3 was used and max target sequences was set to 1, and max hits per sequence was set to 1, resulting in the output of only the top hit. Query sequences were BLASTed against the entire non-redundant database downloaded from the NCBI website on August 2, 2018.

### Gene Ontology Analysis

The program, BLAST2GO provided GO annotations for differentially regulated genes (38) using the following parameters: BLASTx-fast against the nr database, number of blast BLAST hits = 20, E-value of 1.0 e -3, word size of 6, hsp length cutoff of 33, with default mapping and annotation settings. GO terms found to be associated with various genes were manually grouped according to GO subtype (cellular component, biological process, or molecular function) and plotted to view the distribution across time points. The web-based CateGOrizer program was used to batch analyze each set of GO terms and determine the number of GO terms under higher order GO classes of interest (75). WEGO 2.0 (Web Gene Ontology Annotation Plot) with a GO level of 2 was also used to plot histograms of the GO annotations for the differentially regulated genes (45). To further analyze the differentially expressed genes, an enrichment analysis was performed with Metascape (61). A multiple gene list analysis looking at the enrichment of the three classes of Gene Ontology terms was performed using *Drosophila melanogaster* as the analysis species.

## Availability of data and material

Initial description of assembly of this transcriptome in Fisher et al., 2018. Re-assembly was completed as described above and is publicly available at NCBI (Bioproject: PRJNa376023, SUB8325660)

## Declarations

**Ethics approval and consent to participate**

Not applicable.

## Availability of data and materials

Transcriptomic data are available on NCBI (Bioproject: PRJNa376023 (https://www.ncbi.nlm.nih.gov/bioproject/?term=PRJNa376023) and details on the multi-k-mer assembly (GFMG02000000) can be found here: https://www.ncbi.nlm.nih.gov/nuccore/GFMG00000000

## Authors’ contributions

HF, assisted by LL, collected tissue and HF completed the original transcriptome assembly; FW and MM reassembled using multiple K-mers, compacted the new transcriptome using Evigene and did the differential expression analysis; JB, JJ, LMP, and TAM completed the sanger sequencing analysis; DM identified the most stable reference genes; AR and JB completed Protein Yellow phylogenies, LL completed the qPCR, JO and SK consulted on the statistical differential expression analysis; FW wrote the paper; HWH obtained funding for this project and contributed to the writing. All authors read and approved the final manuscript

## Acknowledgements

We thank Marko Melendy for animal care support and Meera Prasad for consulting on revisions to this manuscript.

